# A new mechanism for ribosome rescue can recruit RF1 or RF2 to non-stop ribosomes

**DOI:** 10.1101/465609

**Authors:** Tyler D. P. Goralski, Girish S. Kirimanjeswara, Kenneth C. Keiler

**Author notes:** **Correspondence:** Kenneth Keiler.

## Abstract

Bacterial ribosomes frequently translate to the 3’ end of an mRNA without terminating at an in-frame stop codon. In all bacteria studied to date, these non-stop ribosomes are rescued using *trans*-translation. In some species, genes required for *trans*-translation are essential, but other species can survive without *trans*-translation because they express an alternative ribosome rescue factor, ArfA or ArfB. *Francisella tularensis* cells lacking *trans*-translation are viable, but *F. tularensis* does not encode ArfA or ArfB. Transposon mutagenesis followed by deep sequencing (Tn-seq) identified a new alternative ribosome rescue factor, now named ArfT. *arfT* can be deleted in wild-type cells but not in cells that lack *trans*-translation activity. Over-expression of ArfT suppresses the slow growth phenotype in cells lacking *trans*-translation and counteracts growth arrest caused by *trans*-translation inhibitors, indicating that ArfT rescues non-stop ribosomes *in vivo*. Ribosome rescue assays *in vitro* show that ArfT promotes hydrolysis of peptidyl-tRNA on non-stop ribosomes in conjunction with *F. tularensis* release factors. Unlike ArfA, which requires RF2 for activity, ArfT can function with either RF1 or RF2. Overall, these results indicate that ArfT is a new alternative ribosome rescue factor with a distinct mechanism from ArfA and ArfB.

**Importance:** *Francisella tularensis* is a highly infectious intracellular pathogen that kills more than half of infected humans if left untreated. *F. tularensis* has also been classified as a potential bioterrorism agent with the greatest risk for deliberate misuse. Recently, compounds that inhibit ribosome rescue have been shown to have antibiotic activity against *F. tularensis* and other important pathogens. Like all bacteria that have been studied, *F. tularensis* uses *trans*-translation as the main pathway to rescue stalled ribosomes. However, unlike most bacteria, *F. tularensis* can survive without any of the known factors for ribosome rescue. Our work identifies a *F. tularensis* protein, ArfT, that rescues stalled ribosomes in the absence of *trans*-translation using a new mechanism. These results indicate that ribosome rescue activity is essential in *F. tularensis* and suggest that ribosome rescue activity might be essential in all bacteria.

## Introduction

Bacterial ribosomes frequently translate to the 3’ end of an mRNA that does not have a stop codon (1-3). These non-stop ribosomes cannot terminate translation using one of the canonical termination factors, RF1 or RF2, because they require interactions with the stop codon to activate peptidyl-tRNA hydrolysis (4,5). Data from *E. coli* indicate that 5-10% of ribosomes that initiate translation do not terminate translation at a stop codon on the mRNA, and instead have to be rescued (2,3). The primary ribosome rescue pathway in all bacteria that have been investigated is *trans*-translation (1,2,6). In this pathway, the tmRNA-SmpB complex recognizes a non-stop ribosome and uses a tRNA-like domain of tmRNA and a specialized reading frame within tmRNA to tag the nascent polypeptide for degradation and release the non-stop ribosome (1,2,6-7). Genes encoding tmRNA (*ssrA*) and SmpB (*smpB*) have been identified in >99% of sequenced bacterial genomes, and in some species these genes are essential (1,8). In other species, *trans*-translation is not essential due to the presence of an alternative ribosome rescue factor, ArfA or ArfB (9,10). ArfA is a short protein that inserts its C-terminal tail into the mRNA channel of non-stop ribosomes and rescues them by activating RF2 to hydrolyze the peptidyl-tRNA (10-16). ArfA does not interact with the RF2 residues that recognize a stop codon, but instead binds a different part of RF2 to stabilize the active conformation and promote peptidyl-tRNA hydrolysis (13-16). These interactions cannot be made with RF1, so ArfA only functions in conjunction with RF2 (11-16). ArfB also binds the empty mRNA channel of non-stop ribosomes with its C-terminal tail, but ArfB contains an RF1-like catalytic domain that can hydrolyze peptidyl-tRNA on non-stop ribosomes in the absence of RF1 or RF2 (17-20). In bacteria that have a functional ArfA or ArfB, deletions of *ssrA* and the gene encoding the alternative ribosome rescue factor are synthetically lethal, indicating that these species require at least one mechanism for rescuing non-stop ribosomes (9,10).

Although *ssrA* has been deleted from the pathogen *F. tularensis* (21), no homologs of *arfA* or *arfB* have been found in sequenced *F. tularensis* genomes. *F. tularensis* has a reduced genome size and a life cycle that is different from many other bacteria, so it is possible that ribosome rescue is not essential. Alternatively, *F. tularensis* may have an alternative ribosome rescue system that is sufficiently different from ArfA and ArfB that it cannot be identified by homology searches. *F. tularensis* is a Gram-negative, facultative intracellular bacterium responsible for the vector-borne zoonosis tularemia (22-28). Pneumonic tularemia is infectious at ≤ 10 colony-forming units (cfu) of aerosolized bacteria, and has a 60% mortality rate if left untreated (22-27). *F. tularensis* has been classified as a Tier 1 Select Agent by the CDC because the bacteria can be easily propagated and disseminated as an aerosol, making the threat of a bioterrorist attack with an antibiotic-resistant strain of *F. tularensis* a significant concern (26, 27).

To determine if ribosome rescue is essential in *F*. *tularensis*, we screened for an alternative ribosome rescue factor using transposon mutagenesis followed by deep sequencing (Tn-seq) in *F. tularensis* ssp. holarctica Live Vaccine Strain (LVS). One gene, *FTA_0865*, renamed here as alternative ribosome rescue factor T (ArfT) was found to be essential in cells lacking *trans*-translation but not in wild-type *F. tularensis*. We show that ArfT can rescue non-stop ribosomes *in vivo* and *in vitro*, and that ArfT can function in conjunction with either RF1 or RF2. These data indicate that ribosome rescue is essential in *F. tularensis* and that ArfT is the first representative of a new family of alternative ribosome rescue factors that can recruit either RF1 or RF2 to non-stop ribosomes.

## Results

### Identification of an alternative rescue factor in *F. tularensis*

A published report demonstrated that an *F. tularensis* strain in which *ssrA* was disrupted by insertion of an LtrB intron (*ssrA::LtrB-bp147*) was viable (21). We used RT-PCR to confirm that there was no detectable tmRNA in *ssrA::LtrB-bp147* cells (Fig. S1), suggesting that either ribosome rescue is not essential in *F. tularensis* or that *F. tularensis* has another mechanism to rescue non-stop ribosomes. Homology searches of the *F. tularensis* genome using sequences or motifs from ArfA and ArfB did not identify any candidate alternative ribosome rescue factors. Therefore, we took a genetic approach to identify genes that might be involved in an alternative ribosome rescue pathway. If *F. tularensis* has an unknown alternative ribosome rescue pathway, genes required for the alternative pathway should be essential in *ssrA::LtrB-bp147* cells but not in wild-type cells. We used Tn-seq to identify genes that could be disrupted in each strain. Cells from each strain were mutagenized with a Himar1-based transposon (29, 30) and the transposon insertion sites were sequenced. The ratio of the normalized number of insertions in *ssrA::LtrB-*bp147 to the normalized number of insertions in wild-type was used to identify genes that were much less fit in *ssrA::LtrB-bp147* (Table S1).

Among the genes with no insertions in *ssrA::LtrB-bp147* and typical insertion density in the wild-type strain, *arfT* was a candidate alternative ribosome rescue factor because it shared some characteristics with ArfA and had no annotated function (Fig. 1A). *arfT* encodes a protein of 40 amino acids, whereas mature ArfA has 52-55 amino acids, and ArfT contains a stretch of residues near the C terminus that are similar to a conserved KGKGS sequence found in ArfA (Fig. 1B). Structural studies of ArfA indicate the KGKGS sequence binds in the empty mRNA channel of non-stop ribosomes. A tblastn search (31) showed that ArfT homologues are found in other *F. tularensis* subspecies and in the closely related *F. hispaniensis*, but not in more distantly related species (Table S2). *arfT* was not previously annotated as an open reading frame in *F. tularensis* LVS, the SchuS4 strain, or a number of other sequenced *F. tularensis* strains, but was annotated in *F. tularensis ssp. holarctica* FTNF002-00. For this reason, transposon insertions were mapped to this genome.

**Figure 1:**
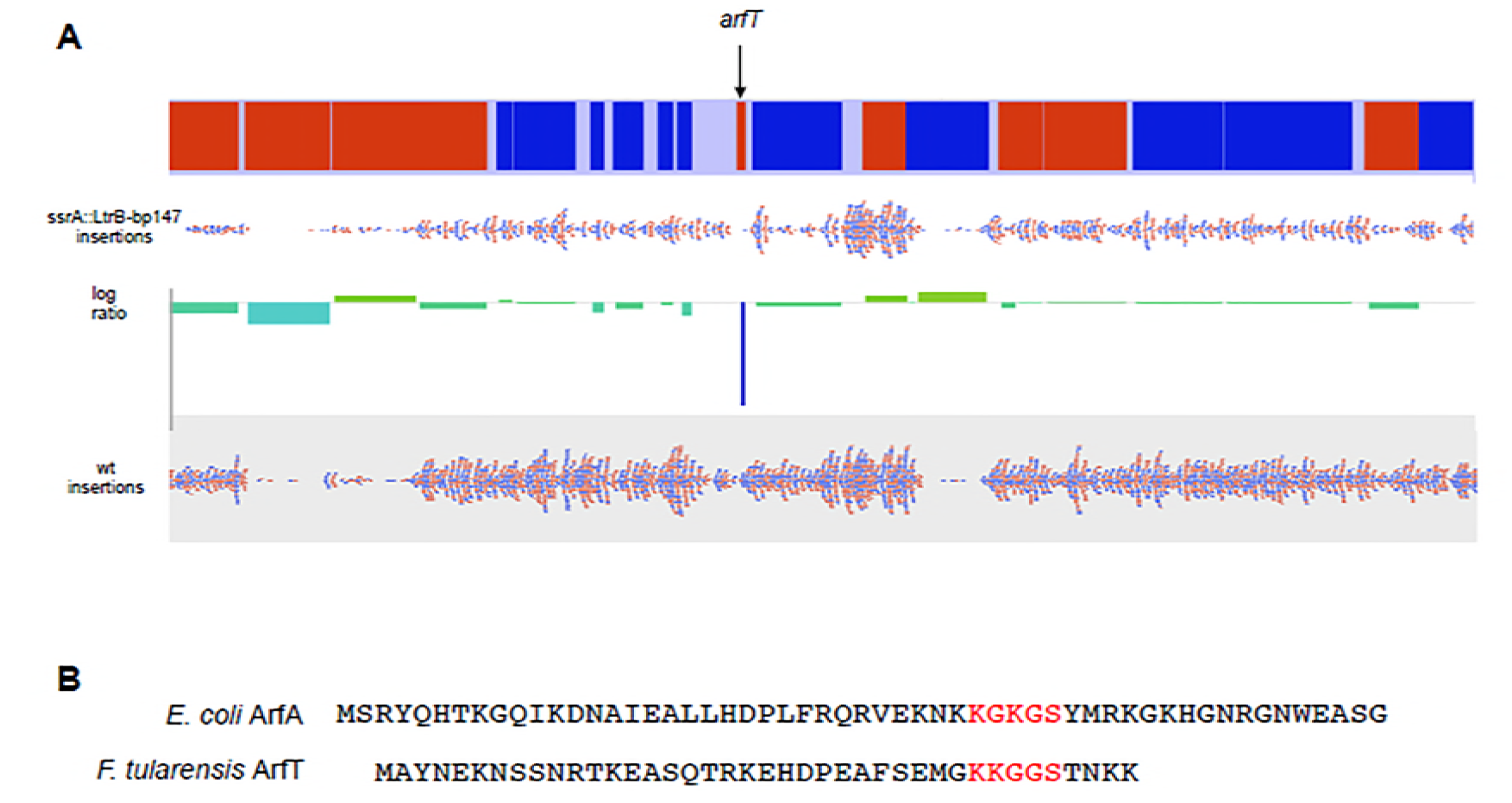
Tn-seq identified arfT as a candidate alternative ribosome rescue system. (A) Representation of Tn-seq data. The portion of the *F. tularensis ssp. holarctica FTNF002-00* chromosome containing *arfT* with genes transcribed to the right in red and those transcribed to the left in blue (top) is shown with mapped transposon insertion sites (red and blue dots) in *ssrA::LtrB-bp147* and wild-type *F. tularensis* (wt). The number of insertions per gene was normalized to the total number of reads and the log ratio of the normalized number of insertions was plotted (center) to identify genes that can be deleted in wild type but not in *ssrA::LtrB-bp147*. (B) Alignment of *E. coli* ArfA and ArfT protein sequences. The KGKGS motif that is conserved in ArfA genes and that binds the empty mRNA channel of the ribosome is shown in red, as are the corresponding residues in ArfT.

### Deletion of *arfT* is synthetically lethal with disruption of *ssrA*

The Tn-seq data suggested that the absence of both *trans*-translation and ArfT is lethal to *F. tularensis* cells. This prediction was tested by attempting to produce markerless, in-frame deletions of *arfT* using a two-step recombination procedure (32) in wild type, *ssrA::LtrB-bp147*, and *ssrA::LtrB-bp147* with a plasmid-borne copy of *ssrA* expressed from a strong, constitutive promoter (*ssrA::LtrB-bp147 pFtssrA*). In the first step of this procedure, a suicide plasmid containing a copy of the *arfT* locus with the *arfT* coding sequence deleted was recombined into the chromosome. The second recombination step eliminates one copy of the *arfT* locus, so cells can retain either the *arfT* deletion or the wild-type *arfT* gene (Fig. S2) (32). The first recombination step was successful in all strains. For the wild-type strain, 20% of the second-step recombinants had the *arfT* deletion, demonstrating that *arfT* is not essential. Deletion of *arfT* did not cause a large defect in growth or morphology (Fig. 3). For the *ssrA::LtrB-bp147* strain, 100 second-step recombinants were screened and all had retained the wild-type copy of *arfT*, indicating that disruption of both *ssrA* and *arfT* was lethal. When a plasmid-borne copy of *ssrA* was present in *ssrA::LtrB-bp147* cells, 20% of the second-step recombinants has *arfT* deleted, demonstrating that the synthetic-lethal phenotype can be complemented by an ectopic copy of *ssrA*. FTA_0993, a gene that had transposon insertions in both wild type and *ssrA::LtrB-bp147* in the Tn-seq experiment, was successfully deleted from the *ssrA::LtrB-bp147* strain (Fig. S2), confirming that *ssrA::LtrB-bp147* cells are competent for recombination in the two-step procedure. Taken together, these data demonstrate that deletion of *arfT* is lethal to *F. tularensis* cells lacking *trans*-translation, and indicate that ribosome rescue is required in *F. tularensis*.

### ArfT can recruit either RF1 or RF2 to hydrolyze peptidyl-tRNA on non-stop ribosomes *in vitro*

*In vitro* ribosome rescue assays were performed to assess whether ArfT was capable of rescuing non-stop ribosomes. Non-stop ribosomes were generated by programming a transcription-translation reaction with a gene that does not have a stop codon (DHFRNS) (Fig. 2) (9). In the absence of ribosome rescue, peptidyl-tRNA is stable on the ribosome and could be observed on protein gels. As expected for non-stop ribosomes, addition of RF1, RF2, and RF3 from *E. coli* or RF1 and RF2 from *F. tularensis* did not dramatically decrease the amount of peptidyl-tRNA. Addition of ArfT alone did not promote hydrolysis of the peptidyl-tRNA, indicating that ArfT does not have intrinsic hydrolytic activity to rescue non-stop ribosomes. Likewise, addition of ArfT in conjunction with RF1, RF2, and RF3 from *E. coli* did not promote peptidyl-tRNA hydrolysis. However, addition of ArfT with *F. tularensis* RF1 resulted in 95% peptidyl-tRNA hydrolysis and addition of ArfT with *F. tularensis* RF2 resulted in 84% peptidyl-tRNA hydrolysis (Fig 2). These data suggest that ArfT can rescue ribosomes by recruiting either RF1 or RF2 to non-stop ribosomes.

**Figure 2:**
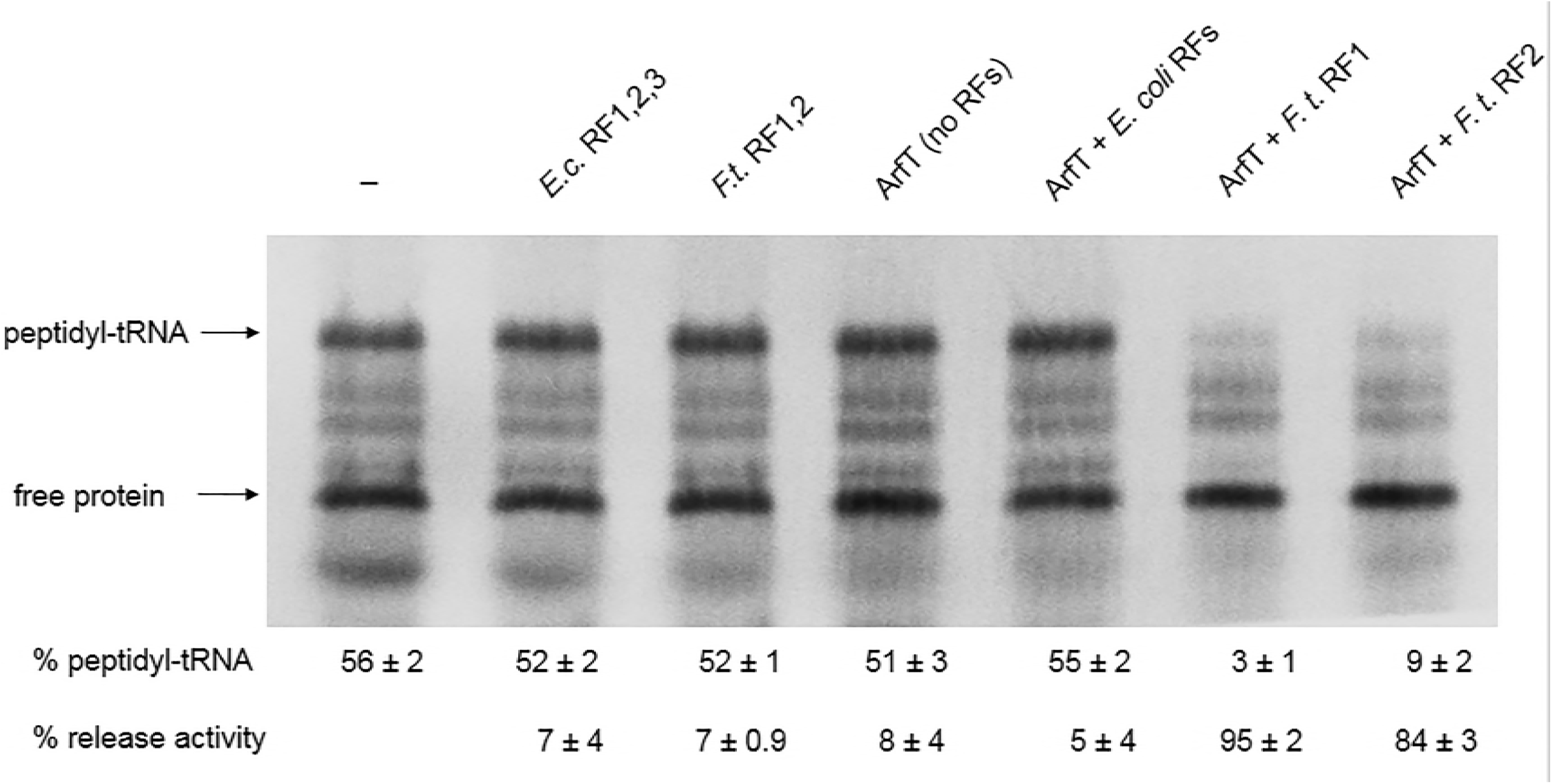
ArfT promotes peptidyl-tRNA hydrolysis on nonstop ribosomes in conjunction with either RF1 or RF2. Gel image of in vitro ribosome rescue assays. In vitro transcription/translation assays were programmed with a non-stop DNA template and synthesized protein was labeled by incorporation of ^35^S-methionine. ArfT and release factors were added to individual reactions in the combinations indicated. Bands corresponding to peptidyl-tRNA and free protein were quantified. The percentage of protein in the peptidyl-tRNA band and the percentage of peptidyl-tRNA that was hydrolyzed compared to a reaction with no release factors or ArfT added (release activity) are shown (± standard deviation). The data are averages of 3 biological replicates.

### Over-expression of *arfT* rescues the growth defect in cells lacking *trans-*translation

It was previously reported that the *ssrA::LtrB-bp147* strain grows much slower than wild type in liquid culture, and that this growth defect could be complemented by expression of *ssrA* from a plasmid (21). To determine whether overexpression of *arfT* could restore normal growth to cells in the absence of *trans*-translation, we cloned *arfT* under the control of the strong, constitutive bacterioferratin (Bfr) promoter on a multicopy plasmid (pArfT) and tested its impact on growth rate. As expected, the *ssrA::LtrB-bp147* grew substantially slower than wild-type, but *ssrA::LtrB-bp147 pFtssrA* grew at the same rate as wild-type (Fig. 3). *ssrA::LtrB-bp147* cells with pArfT also grew at the same rate as wild-type, indicating that multi-copy *arfT* can suppress the *ssrA* phenotype. pArfT did not increase the growth rate of wild-type cells (Fig. 4). These results suggest that ArfT can rescue non-stop ribosomes in the absence of *trans*-translation.

**Figure 3:**
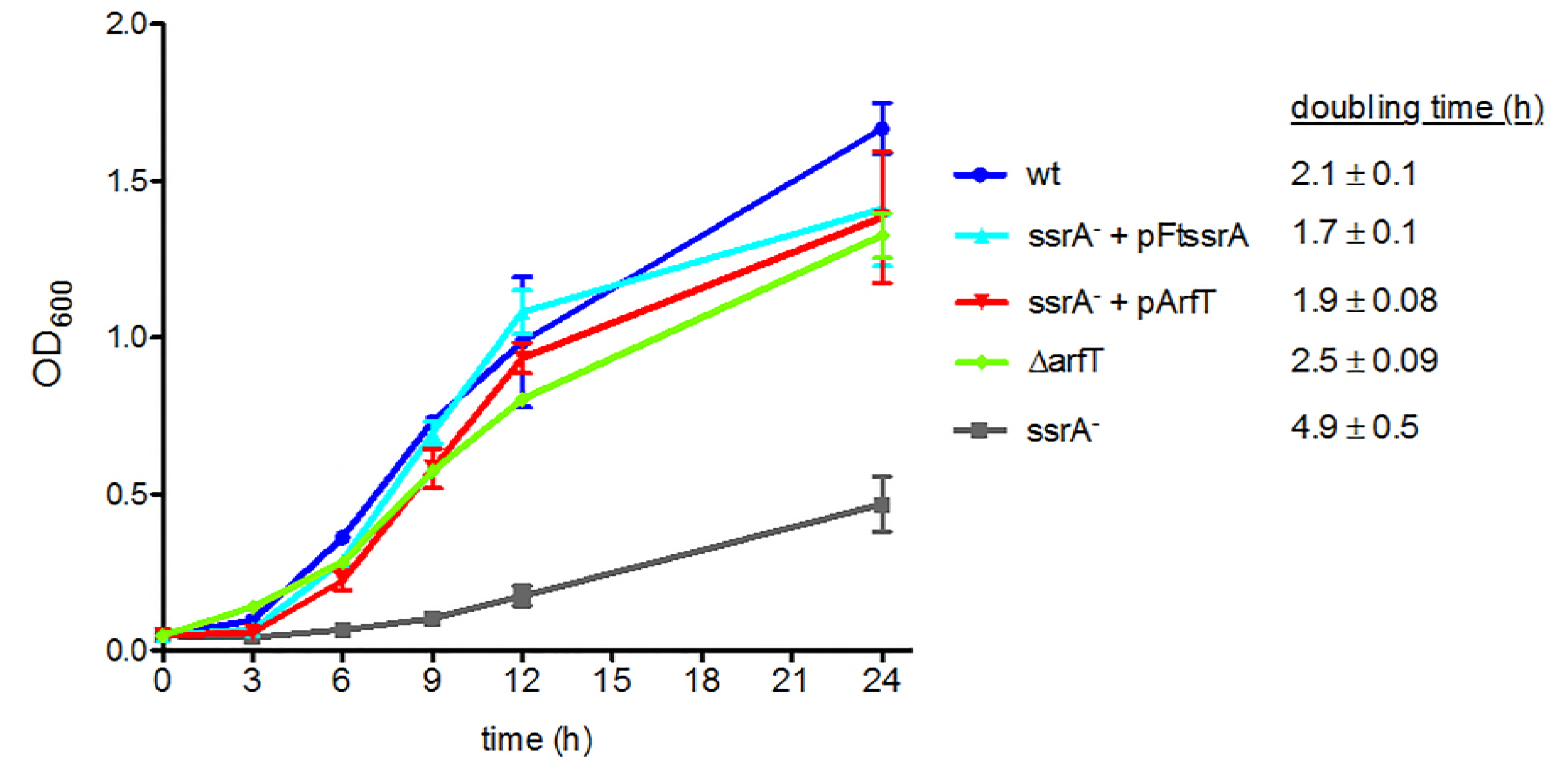
Overexpression of ArfT rescues the growth defect in *ssrA::LtrB-bp147*. Growth curves of wild-type *F. tularensis* (wt), the ∆*arfT* strain, and the *ssrA::LtrB-bp147* strain (*ssrA^−^*) with and without plasmids expressing *ssrA* (pFtssrA) or *arfT* (pArfT). Error bars indicate standard deviation. The doubling time for each strain (± standard deviation) is indicated. The data are averages of 3 biological replicates.

**Figure 4:**
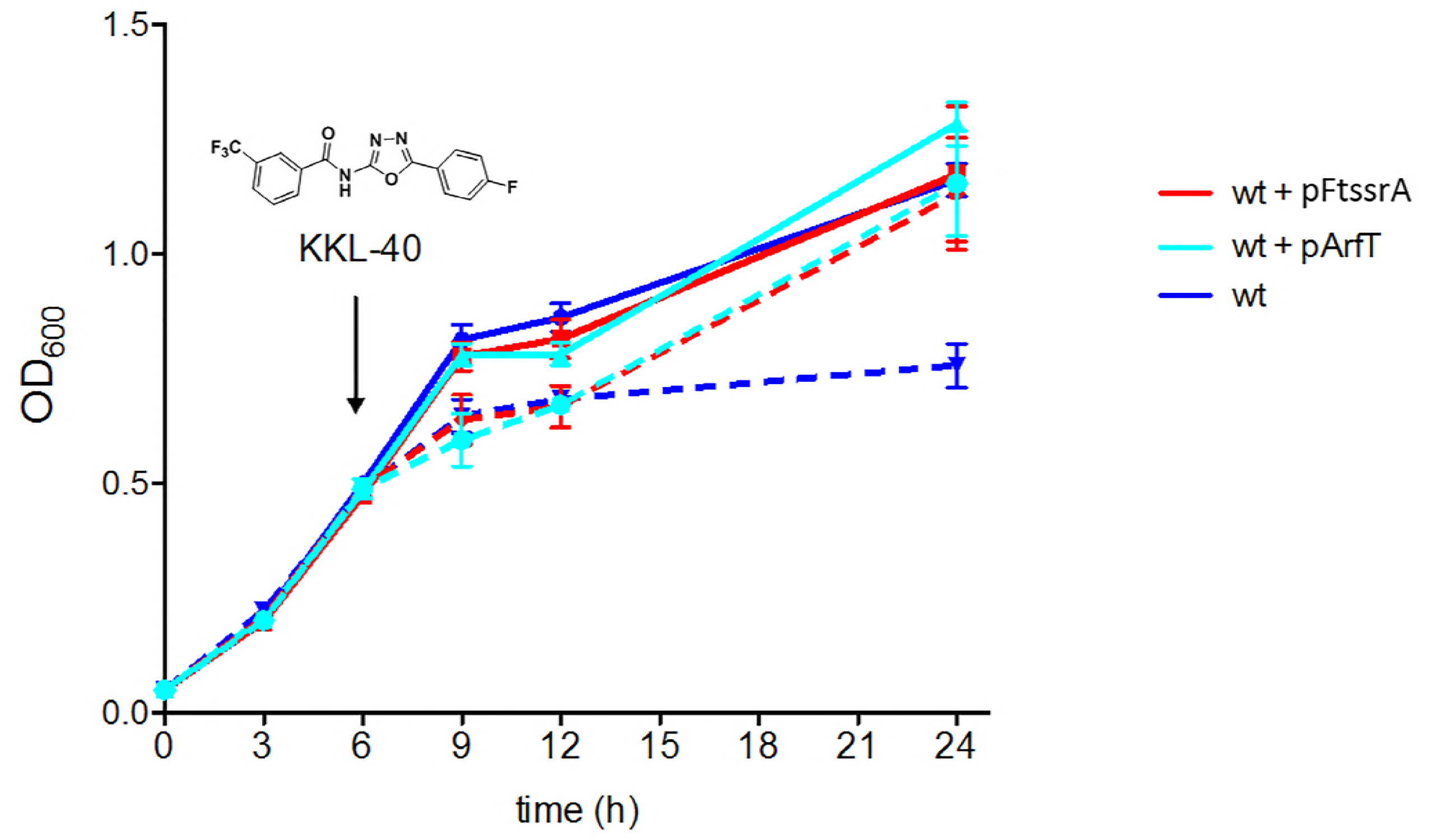
Over-expression of ArfT prevents growth inhibition caused by ribosome rescue inhibitors. Growth curves of wild-type *F. tularensis* (wt) with and without plasmids expressing *ssrA* (pFtssrA) or *arfT* (pArfT). A ribosome rescue inhibitor, KKL-40 (structure shown), was added to half the cultures after 6 h (indicated by arrow) at 10X MIC. Cultures with KKL-40 are shown by dotted lines and cultures with no drug are shown by solid lines. The data are averages of 3 biological replicates, with error bars indicating the standard deviation.

### Over-expression of ArfT prevents growth arrest due to ribosome rescue inhibitors

It has been shown that a class of oxadiazole compounds such as KKL-40 inhibit ribosome rescue and arrest the growth of many bacterial species, including *F. tularensis* (33-35). Over-expression of *E. coli* ArfA prevents growth arrest by these oxadiazoles in *Shigella flexneri*, confirming that growth arrest is due to inhibition of ribosome rescue (33,34). If ArfT has ribosome rescue activity similar to ArfA, over-expression of ArfT should inhibit growth arrest in *F. tularensis* by KKL-40. To test this prediction, KKL-40 was added to growing cultures of *F. tularensis* strains and growth was monitored over 18 h (Fig. 4). As previously observed, addition of KKL-40 resulted in growth arrest of wild-type *F. tularensis* and the bacteria were unable to recover to normal levels. Addition of KKL-40 to *F. tularensis* carrying pFtssrA or pArfT caused an initial decrease in growth rate, but after 18 h the cultures had reached the same density as wild type. Because growth inhibition is suppressed by extra ribosome rescue activity in the form of either tmRNA-SmpB or ArfT, it is likely that KKL-40 inhibits growth through ribosome rescue and not through off-target effects. These results are consistent with a model in which KKL-40 arrests growth in *F. tularensis* by binding to non-stop ribosomes and tmRNA-SmpB or ArfT can counteract the effects of KKL-40 by rescuing the ribosomes before KKL-40 binds.

## Discussion

The data described here answer two of the recently posed outstanding questions for ribosome rescue: Are there other alternative rescue factor systems, and will ArfA-like systems emerge in bacteria where RF1 is recruited to non-stop ribosomes (36)? The answer to both questions is yes. The data presented here indicate that ArfT has all the characteristics of an alternative ribosome rescue factor in *F. tularensis*. ArfT has ribosome rescue activity *in vitro* because it can release non-stop ribosomes in conjunction with RF1 or RF2. *In vivo*, deletion of *arfT* is synthetically lethal with disruption of *ssrA*, consistent with ArfT providing essential ribosome rescue activity in the absence of *trans*-translation. Over-expression of ArfT suppresses the slow growth phenotype in cells lacking *trans*-translation and counteracts growth arrest by a ribosome rescue inhibitor in *F. tularensis*, indicating that ArfT can perform the same physiological role as *trans*-translation in *F. tularensis*. These results demonstrate that the presence of ArfT in *F. tularensis* makes *trans*-translation dispensable, and that ribosome rescue activity is essential in *F. tularensis*.

ArfT has some similarities to ArfA and these factors may recognize non-stop ribosomes in the same manner. The C-terminal tail of ArfA binds in the empty mRNA channel of non-stop ribosomes through a number of lysine and arginine residues including a conserved KGKGS motif (13-16). None of these residues are essential for ArfA activity (16, 37), but replacement of individual residues decreases ribosome rescue activity *in vitro* (16). The KKGGSTNKK sequence near the C-terminus of ArfT has an arrangement of positively charged residues that is similar to those in ArfA, suggesting that ArfT may use this sequence to bind the ribosome. SmpB and ArfB also bind in the empty mRNA channel of non-stop ribosomes using positively charged C-terminal tails, but ArfA, SmpB, and ArfB each make different interactions with the mRNA channel (7,13-20,37). Because of this variation in binding, structural studies will be required to define the interactions between ArfT and the ribosome.

Despite the similarities in protein size and C-terminal tail sequence between ArfT and ArfA, the observation that ArfT can activate RF1 or RF2 suggests that ArfT may not interact with release factors in the same way as ArfA. Cryo-EM analyses of a non-stop ribosome bound to *E. coli* ArfA-RF2 showed that residues 15-31 of ArfA interact with RF2 to stabilize the active conformation of RF2 and promote hydrolysis of the peptidyl-tRNA (13-16). In a key feature of this interaction, ArfA forms a ß-strand that extends the ß-sheet formed by ß4-ß5 of RF2, with F25 of ArfA binding in a hydrophobic pocket formed by V198 and F217 of RF2. Residues in RF2 ß4-ß5 and the SPF loop are highly conserved between *E. coli* RF2 and *F. tularensis* RF2 (Fig. S3), raising the possibility that ArfT could bind in a similar manner as ArfA. However, ArfT does not have a hydrophobic residue at the position corresponding to F25 (Fig. 1B). The absence of the V198-F217 pocket in *E. coli* RF1 has been suggested to be the reason ArfA does not activate *E. coli* RF1 (13-16). This region of *E. coli* RF1 is highly conserved in *F. tularensis* RF1, yet ArfT activates *F. tularensis* RF1 but not *E. coli* RF1. Therefore, if the interaction between ArfT and RF2 is similar to the interaction between ArfA and RF1, ArfT would have to activate RF1 through a distinct mechanism. Alternatively, ArfT may activate *F. tularensis* RF1 and RF2 in the same manner, but through a different mechanism than that used by ArfA. Little was known about the interactions among ArfA, RF2, and the ribosome before structural data of the complex was obtained, and similar studies will be required to understand how ArfT can activate both RF1 and RF2.

Another likely difference between ArfT and ArfA is regulation. The *arfA* gene includes a transcriptional terminator and RNase III cleavage site before the stop codon, such that ArfA protein is made from non-stop mRNA (38,39). When *trans*-translation is active, the nascent ArfA peptide is tagged and degraded, but if *trans*-translation activity is not available active ArfA is produced and accumulates in the cell. This genetic arrangement makes ArfA a true backup ribosome rescue system, functioning only when *trans-*translation activity is low or absent (38,39). The *arfT* gene does not include a transcriptional terminator or an RNase III cleavage site before the stop codon. RT-PCR using a primer corresponding to the final 33 nucleotides of the *arfT* reading frame (including the stop codon) showed that *arfT* mRNA accumulated in wild-type *F. tularensis* and the *ssrA*-disrupted strain at similar levels (Fig. S4). Although these results do not exclude the possibility that *arfT* mRNA is truncated in the last few codons, it does not appear to be controlled by transcriptional termination and RNase III cleavage in the same manner as ArfA.

The observations that ArfT interacts with RF1 and is not regulated like ArfA, and the overall low sequence similarity between ArfT and ArfA, suggest that ArfT evolved independently from ArfA and represents a third different alternative ribosome rescue factor. Our sequence homology searches only identified ArfT in closely related *F. tularensis* and *F. hispaniensis* strains, but the small size of ArfT makes more distant homologs difficult to identify with this method. Characterization of the ArfT residues required for interaction with RF1 and RF2 will allow more specific searches for ArfT in other species. The number of different ribosome rescue mechanisms discovered to-date suggests that the problem presented by non-stop ribosomes has been solved many times throughout evolution, and more alternative ribosome rescue factors may yet be discovered. It is not yet clear what conditions would limit *trans*-translation activity enough that an alternative ribosome rescue factor would be needed. However, such conditions must exist in a wide variety of environments. Alternative ribosome rescue factors have been selected for in enteric bacteria such as *E. coli*, which has ArfA, aquatic bacteria such as *C. crescentus*, which has ArfB, and intracellular pathogens such as *F. tularensis*, which has ArfT.

## Materials and Methods

### Bacterial culture

Bacterial strains are listed in Table 1. *E. coli* DH10B was used for routine cloning procedures and was grown in Luria–Bertani (LB) broth (10% bacto-tryptone, 5% yeast extract, 10% NaCl, [pH 7.5]), or on LB agar supplemented with ampicillin (100 μg/mL), or kanamycin (30 μg/mL) where appropriate. *F. tularensis* was grown in Chamberlain’s defined medium (CDM) (40) adjusted to pH 6.2 at 37°C with shaking, or on chocolate agar plates (Mueller-Hinton agar supplemented with 1% bovine hemoglobin [Remel, USA] and 1% Isovitalex X Enrichment [Becton Dickinson, France]) at 37°C in a humidified incubator with 5% CO_2_ for 48-72 h. Kanamycin (10 μg/mL), tetracycline (10 μg/mL), and sucrose (5%) were added to cultures and plates where appropriate. For growth curve experiments, *F. tularensis* cultures were grown in CDM overnight at 37°C and 200 rpm and back diluted to an optical density at 600 nm (OD_600_) of 0.05. Growth was monitored by performing OD_600_ readings. When indicated, 1.4 μg/mL KKL-40 was added 6 h post inoculum.

### Plasmid construction

Oligonucleotide sequences are provided in Table S3 in the supplemental material. To generate plasmids pMP812-ΔArfT and pMP812-Δ0993 600 basepair PCR products flanking the gene to be deleted were amplified using primers ArfT_UF, ArfT_UR, ArfT_DF, ArfT_DR, and 0993_UF, 0993_UR, 0993_DF, 0993_DR, digested with BamHI, and ligated together. The sequence was then reamplified as one unit with primers ArfT_UF, ArfT_DR, and 0993_UF, 0993_DR, and cloned into pMP812 using SalI and NotI restriction sites. Plasmids pArfT and pFtssrA were constructed by amplifying the coding sequences of each gene using primers ArfT_CF and ArfT_CR, and FtssrA_CF and FtssrA_CR. The Bfr promoter (41) was amplified using primers Bfr_F and Bfr_R, ligated upstream of either the ArfT or ssrA PCR product using a BamHI restricition site, and reamplified as one unit with primers Bfr_F, and either ArfT_CR or ssrA_CR. The resulting PCR product was digested with Eco RI and ligated into the plasmid pKK214-MCS_4_ (41). In order to construct plasmids pET28ArfT, pET28RF1, and pET28RF2, primers RF1_PF, RF1_PR, RF2_PF, RF2_PR, and ArfT_PF, ArfT_PR, were used to generate PCR products of the protein coding sequence of RF1, RF2, and ArfT from *F. tularensis*. The PCR products were then cloned into pET28a(+) using NdeI and XhoI restriction sites for protein expression of ArfT, as well as release factor 1 (RF1), and release factor 2 (RF2) from *F. tularensis*.

### Tn-seq

Overnight cultures of wild-type *F. tularensis* and the *ssrA::LtrB-bp147* strain were grown to OD_600_ = 0.5, washed 3x with 500 mM sucrose, and transformed with ~300 ng of the plasmid pHimar H3. Over 50,000 colonies were pooled, and chromosomal DNA was extracted. The libraries were prepared and sequenced on an Illumina HiSeq 2000 by Fasteris (location). The data were mapped to the genome of *F. tularensis ssp. holarctica* FTNF002-00, and analyzed in Geneious version 11.1.4 using parameters described previously (9). The frequency of transposition for each gene was quantified in both strain backgrounds. Additionally, the relative fitness of each gene in both strains was quantified by looking at the ratio of the number of times a sequence was recovered in the *ssrA* mutant as compared to wt. Insertion ratio data was generated for each gene to determine if genes were essential in the absence of ssrA (Table S1).

### Purification of ArfT, *F. tulanesis* RF1, and *F. tulanesis* RF2

Strains TG001, TG002, and TG003 were grown to OD_600_ ~ 0.8, and the expression of ArfT, RF1 or RF2 was induced by the addition of isopropyl-β-D-thiogalactopyranoside (IPTG) to 1 mM. Cells were harvested by centrifugation, resuspended in native lysis buffer (50 mM sodium phosphate 300 mM NaCl, 5 mM imidazole, [pH 8.0]), and sonicated or processed through a French press. The lysate was cleared by centrifugation at 14,000 *g* for 10 min. Ni-nitrilotriacetic acid (NTA) agarose (Qiagen) that had been equilibrated with lysis buffer, was added to the cleared lysate, followed by incubation with gentle rocking at 4 °C for 1 h. The slurry was packed in a column, and washed with 10 volumes of native wash buffer (50 mM sodium phosphate, 300 mM NaCl, 20 mM imidazole, [pH 8.0]). Bound protein was eluted with native elution buffer (50 mM sodium phosphate, 300 mM NaCl, 250 mM imidizole, [pH 8.0]), and visualized by SDS-PAGE. Fractions containing 6×His-protein were dialyzed against RF or ArfT storage buffer (50 mM HEPES, 300 mM NaCl, [pH 7.5] for FTA_0865, 50 mM Tris-HCl, 300 mM NaCl, [pH 7.0] for RF1 and RF2). The 6xHis-tag was removed from RF1 with the Thrombin CleanCleave Kit (Sigma-Aldrich) following the manufacturer’s instructions. The cleaved RF1 protein solution was loaded with NTA agarose, incubated with gentle rocking at 4 °C for 1 h. The slurry was packed into a column, and the flow-through containing RF1 was collected. RF2 was dialyzed against Buffer A (50 mM Tris-HCl, 100mM NaCl, [pH 7.0] and purified on a MonoQ column using an AKTA purifier (GE Healthcare Life Sciences). Proteins were visualized by SDS-PAGE and dialyzed into RF storage buffer.

### *In vitro* translation and peptidyl hydrolysis assays

ArfT peptidyl hydrolysis activity was assessed using a previously described assay (9). Briefly, non-stop DHFR was PCR amplified with primers HAF_T7 and UTR_DHFR_FL, added to the PURExpress ΔRFs system (New England Biolabs) A and B reaction mixtures and incubated for 1 h at 37°C. Where indicated, ArfT was added to a final concentration of 25 μg/mL and *E. coli* or *F. tularensis* LVS RFs were added to a final concentration of 500 μg/mL, and the reactions were incubated for 1 h at 37°C. Total protein was precipitated by addition of cold acetone, resuspended in sample loading buffer (5 mM sodium bisulfite, 50 mM MOPS [morpholinepropanesulfonic acid], 50 mM Tris base, 1 µM EDTA, 0.1% SDS, 5% glycerol, 0.01% xylene cyanol, 0.01% bromophenol blue), and resolved on a Bis-Tris gel using MOPS running buffer.

### Genetic deletions

Targeted, markerless in-frame deletions were generated for both *FTA_0865* and *FTA_0993* with a two-step allelic exchange system designed for *F. tularensis* using the pMP812 *sacB* suicide vector (32). *F. tularensis* strains were transformed with either pMP812-ArfT or pMP812-0993, and primary recombinants were selected on kanamycin after incubating at 37°C in a humidified incubator with 5% CO_2_ for 48-72 h. Primary recombinants were grown overnight without selection and plated on 5% sucrose to select for secondary recombinants. Secondary recombinants were confirmed by replica plating on chocolate agar containing kanamycin, and chocolate agar without selection. Genetic deletions were confirmed via PCR using primers ArfT_KOF and ArfT_KOR, and 0993_KOF and 0993_KOR.

## Acknowledgments

The authors thank Wali Karzai for his contribution of the *ssrA::LtrB-bp147 F. tularensis* strain, and Dara Frank for her contribution of the Himar H3 plasmid. This work was funded by the Science Mathematics and Research for Transformation (SMART) Scholarship (to T.G.) and National Institutes of Health Grant GM121650 (to K.C.K.).

## Supplemental Figure Legends

**Figure S1. No tmRNA is present in *ssrA::LtrB-bp147* cells.** qPCR results for amplification of tmRNA from cDNA prepared from wild-type *F. tularensis* (wt) or the *ssrA::LtrB-bp147* strain. The relative florescence units (RFU) are plotted as a function of PCR cycle, and the positive amplification threshold is indicated by the black line. These results show the amount of tmRNA in *ssrA::LtrB-bp147* is decreased by a factor of >10^8^ compared to the amount in wt.

**Figure S2. Deletion of *arfT* is synthetically lethal with deletion of *ssrA***. Diagnostic PCR was performed to determine if secondary recombinants from allelic exchange were deletions or reversions to wild type. One example of each strain is shown. Lanes 2-5 are PCR products from reactions using ArfT_KO primers, and lanes 6-9 are PCR products from reactions using 0993_KO primers. The ArfT_KO primers amplify a 690 bp product for the wild type and a 570 bp product for a deletion of *arfT*. The 0993_KO primers amplify a 670 bp product for the wild type and a 330 bp product for deletions. Lanes: 1. DNA marker; 2. wt control DNA; 3. *arfT* deletion in wt; 4. *arfT* deletion in *ssrA::LtrB-bp147* + pFtssrA; 5. reversion to wild type in *ssrA::LtrB-bp147*; 6. Wt control DNA; 7. FTA_0993 deletion in wt; 8. FTA_0993 deletion in *ssrA::LtrB-bp147* + pFtssrA; 9. FTA_0993 deletion in *ssrA::LtrB-bp147*.

**Figure S3. Residues in the ArfA-interacting region of *E. coli* RF2 are conserved in *F. tularensis* RF2.** Alignments of RF2 from *E. coli* and *F. tularensis* using blastp (26). Residues in RF2 ß4 (red), ß5 (blue), and the SPF loop (gold) are indicated.

**Figure S4. *arfT* mRNA accumulates in *F. tularensis*.** cDNA from wild type (wt), *ssrA::LtrB-bp147*, wild expressing *arfT* from a plasmid (wt + pArfT), or wild type expressing *ssrA* from a plasmid (wt + pFtssrA) was used as template for qPCR amplification of *arfT*. The relative florescence units (RFU) are plotted as a function of PCR cycle, and the positive amplification threshold is indicated by the black line.

